# Predicting Future Kinetic States of Physicochemical Systems Using Generative Pre-trained Transformer

**DOI:** 10.1101/2024.05.22.595440

**Authors:** Palash Bera, Jagannath Mondal

## Abstract

Capturing the time evolution and predicting future kinetic states of physicochemical systems present significant challenges due to the precision and computational effort required. In this study, we demonstrate that the transformer, a machine learning model renowned for machine translation and natural language processing, can be effectively adapted to predict the dynamical state-to-state transition kinetics of biologically relevant physicochemical systems. Specifically, by using sequences of time-discretized states from Molecular Dynamics (MD) simulation trajectories as input, we show that a transformer can learn the complex syntactic and semantic relationships within the trajectory. This enables this generative pre-trained transformer (GPT) to predict kinetically accurate sequences of future states for a diverse set of models and biomolecules of varying complexity. Remarkably, the GPT can predict future states much faster than traditional MD simulations. We show that it is particularly adept at forecasting the time evolution of an out-of-equilibrium active system that do not maintain detailed balance. An analysis of self-attention mechanism inherent in transformers is found to hold crucial role for capturing the long-range correlations necessary for accurate state-to-state transition predictions. Together, our results highlight the ability of transformer based machine learning model in generating future states of physicochemical systems with statistical precision.

## Introduction

The time evolution of any physical system undergoes various state-to-state transitions. Understanding the dynamics of these systems particularly at the molecular level, poses a significant challenge due to the complexity of their transitions between various states. Traditional approaches, such as molecular dynamics simulations (MDs) offer valuable insights into these transitions. However, the MDs are computationally very expensive, limiting their applicability to large-scale systems or long-time predictions. Moreover, describing the actual phase space of a physical system typically involves handling data of very high dimensions. The lower dimensional representation of this data along some order parameters can provide information about various states and transitions between them. Nonetheless, achieving a comprehensive understanding of these transitions over a very long time requires performing highly resource-intensive MD simulations.

In recent years, state-of-the-art machine learning (ML)/artificial intelligence (AI), especially recurrent neural networks (RNN) and large language models (LLM) have become a promising tool in addressing these challenges.^1–7^ The RNN-based ML architecture was developed for time series prediction, particularly for forecasting the weather, electricity consumption, revenue, etc.^8–11^ They are mainly known for their ability to remember the memory of past inputs while processing current ones. Their capability to capture sequential patterns makes them particularly well-suited for tasks where the order of data matters. However, RNNs suffer from a key limitation known as the vanishing gradients problem.^12^ As the length of the input sequence increases, the gradients used to update the RNN’s weights tend to become smaller and smaller. To address this issue, an advanced version of RNNs called Long Short-Term Memory (LSTM)^13^ was developed. LSTM can solve this vanishing gradients problem by using specialized memory cells and a gating mechanism. They can dynamically adjust their memory by remembering or forgetting information across multiple time steps. This enables them to effectively capture long-term patterns present in sequential data.

Apart from the forecasting problem, the LSTM-based architecture can be used for a variety of sequence-to-sequence tasks, such as language modelling, machine translation, and speech recognition.^2,14,15^ However, for large language modelling (LLM) the LSTM might not work to capture the complex syntactic and semantic relationships present in natural language. To overcome these issues, the pioneering work by Vaswani et al.^7^ introduced the attention-based model known as Transformer. The concept of self-attention mechanisms can encode contextual information from the input text and generate coherent and contextually relevant responses. Although the original work of the Transformer was mainly designed for machine translation, the various parts of the Transformer can be used for different purposes. For instance, the encoder component can be applied to classification problems, while the decoder component can be used for sequence-to-sequence generation.

In our study, we have utilized the decoder component of the Transformer architecture to predict the future kinetic states of diverse physicochemical as well as biologically relevant systems. Our investigations primarily focus on set of systems of hiererchical complexity: ranging from hand-crafted three-state and four-state model potentials, a globular folded protein namely Trp-cage to an intrinsically disordered protein (IDP) *α*-Synuclein. We demonstrate that our protocol can effectively learn the time evolution of different states in MD or other kinetic simulations along certain low-dimensional order parameters. As a result, the model can generate future states that are kinetically and thermodynamically accurate. Interestingly, the model is remarkably powerful, as it can accurately generate the future state even for an active system that is out of equilibrium, as would be demonstrated for a 32-bead active worm-like polymer chain and its passive counterpart. Moreover, for the more complex systems, we have found that the attention mechanism plays a crucial role in maintaining the relationships between the states, enabling the model to generate the future state correctly.

## Results

### Learning Molecular dynamics trajectory using transformer model

In any language model, the input is a series of characters or words. During the training process, the weights and biases of the various layers are optimized, enabling the model to understand the context and relationships between different parts of the input sequence. Once the model is trained, it can generate the subsequent sequence for a given input sequence in an auto-regressive manner. As illustrated in figure 1, here we implement a comprehensive scheme to learn kinetic trajectory via transformer and then to generate future states that could be segmented into three stages:

**Figure 1.**
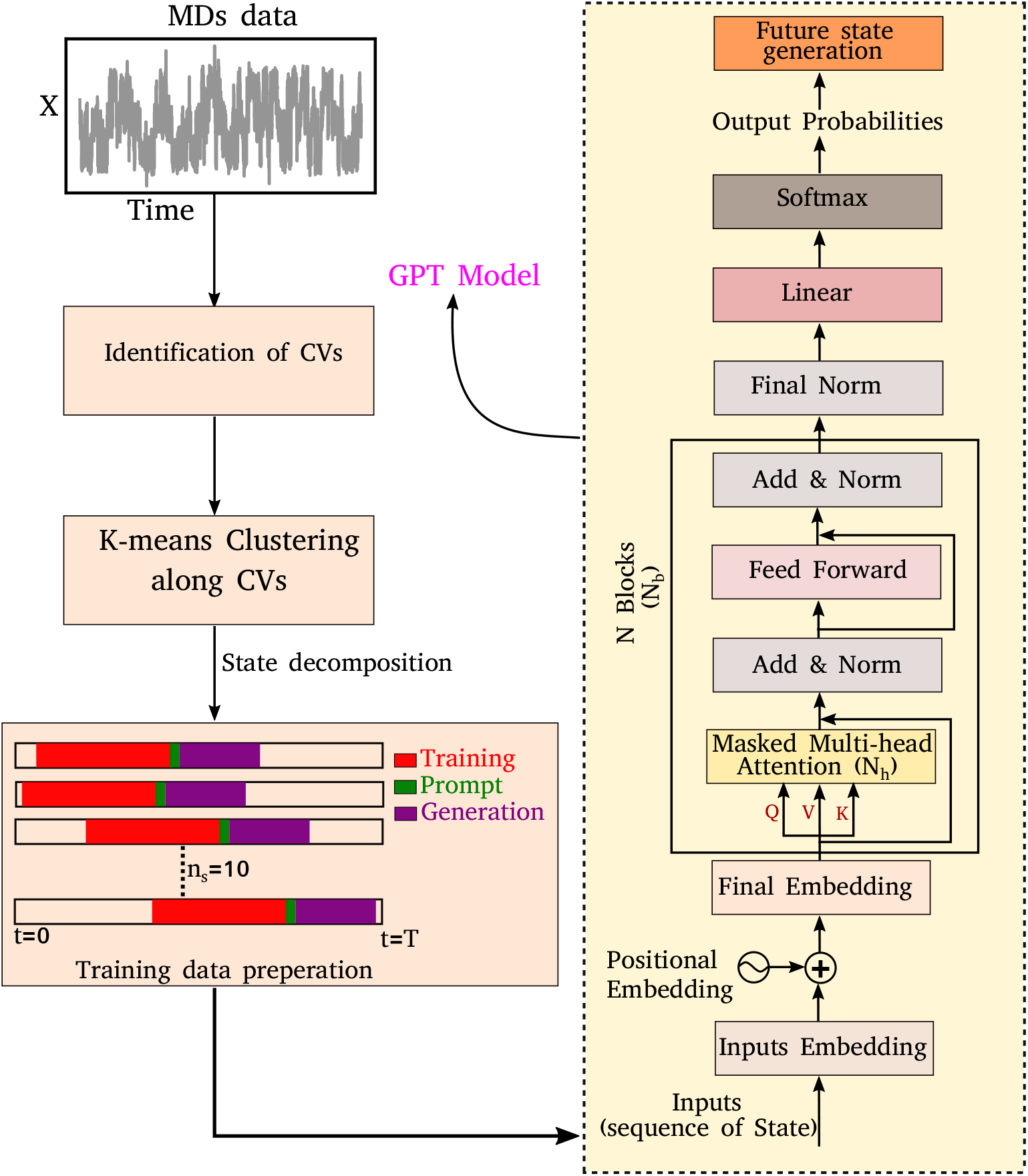
A schematic representation of training data preparation and the architecture of the decoder-only transformer. The left-hand side figure represents the schematic representation of the discretization of molecular dynamics simulation (MDs) trajectory achieved through the identification of collective variables (CVs) and K-means clustering. A total of *n*_*s*_ = 10 segments are randomly selected from the discretized trajectory to train an equal number of independent generative pre-trained transformer (GPT) models. Each trained model generates subsequent sequences starting from where the respective segment ended, using a few sequences as prompts. The right-hand side figure depicts the various layers of a decoder-only transformer. The model architecture consists of input embedding, positional embeddings, and multiple blocks of masked multi-head attention, normalization, and feed-forward layers. The model is optimized using cross-entropy as the loss function.

- A. Segmentation of MD trajectory into discrete states.
- B. Training a decoder-only transformer using MD-derived states as input.
- C. Generating future states from pre-trained transformer.

Below we describe the different stages of our scheme.

#### A. Discretization of Molecular Dynamics trajectory into meaningful state-space

The time evolution of a physical system involves various conformational or state changes and transitions between them. Our study aims to predict the future state of physicochemical systems using a large language model, specifically a decoder-only transformer. Interestingly, the state prediction problem can be mapped to the sequence generation of any language model where each state corresponds to a distinct vocabulary item within a corpus. For all of the systems, we have utilized molecular dynamics (MD) simulation trajectories to predict their future state. However, MD trajectories are continuous, requiring discretization of the trajectory into grid points or states before inputting it into a language model. For simple systems, particle position can be used for discretization. However, complex systems with high dimensionality or degrees of freedom require defining some order parameters or collective variables (CVs) to discretize the trajectory into a certain number of states. To identify the various states of a physical system, we performed K-means clustering along CVs. Consequently, the trajectory is segmented into distinct sequences of states. These sequences can then be used as inputs for a decoder-only transformer model. Hereafter, these sequences of state will be referred to as sequences of token. For training the decoder-only transformer model, we randomly chose *n*_*s*_ number of segments from the discretized trajectory and trained *n*_*s*_ number of independent models with the same architecture. From each independent trained model, we generated the next sequences from where the corresponding segment ended by providing a few sequences as a prompt. All the results presented here are averaged over these independent runs. The left side of Figure 1 illustrates a schematic overview of this process. The specific details regarding the amounts of data used for training, validation, and testing across the various systems are provided in Table-S1. Our approach and model are different from the methodology employed by Tsai et al.^16,17^ In their study, they utilized an LSTM-based architecture to learn the MD simulation trajectory, and the states were generated from the training data itself.

#### B. Training a decoder-only transformer with MD-derived states as input

Now we will delve into the architecture of the decoder-only transformer model. The neural network-based decoder-only transformer^7^ consists of several layers as depicted in the right side of the Figure 1. The first layer is an input embedding or token embedding layer which transforms each token of the input sequence into a dense vector. The dimension of the vector is a hyperparameter which is called embedding dimension. For a sequence of length *l* and embedding dimension *d*, this layer transforms each token into a d-dimensional vector. Consequently, the embedding layer generates a (*l*×*d*)-dimensional matrix, often referred to as the embedding or weight matrix. Throughout the training process, the elements of this weight matrix undergo optimization and the model will learn the semantic relationship between different tokens of the time sequence data.

To enable the GPT model to learn the sequence order of time series data, the positional information for each token is required which can be achieved through positional embeddings. For an input sequence of length *l*, the position of the *k*^*th*^ token can be represented as follows^7^

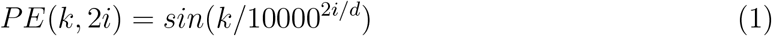

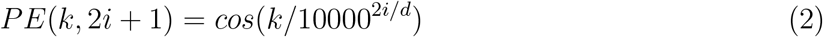

where *d* is the dimension of output embedding and, for each *k*(0 ≤ *k* < *l*), *i* can take value from 0 to *d/*2. Hence, one can achieve the final embedding layer by adding these two embeddings.

The final embedding layer is followed by multiple blocks of layer (*N*_*b*_). Each block typically comprises various components, including masked multi-head attention with a specified number of heads, *N*_*h*_, normalization layers, and feed-forward layers. Among these, the masked multi-head attention layer is particularly significant, serving as a communication mechanism between each token in the sequence. To calculate the key (K), query (Q), and value (V) tensors, the final embedded vector, denoted as *X*_*f*_, is multiplied with three trainable weight matrices (*W*_*k*_, *W*_*q*_, and *W*_*v*_) as: *K* = *X*_*f*_ *· W*_*k*_, *Q* = *X*_*f*_ *· W*_*q*_, and *V* = *X*_*f*_ *· W*_*v*_.

The attention score *A*_*s*_ is then calculated as 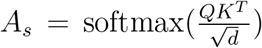, and the output of the attention layer is given by^7^

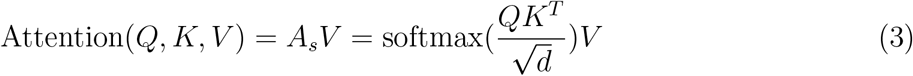

where softmax 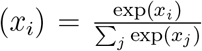. This mechanism enables the model to discern the relative importance of tokens to each other, providing a clear sense of context and relationships between them.

Finally, the normalized outputs of the *N*_*b*_ layer are used as inputs of a fully connected dense linear layer. Given that the transformer model functions as a probabilistic model, the output of this dense layer is passed through a softmax function to yield categorical probabilities. We have used cross-entropy as our loss function to minimize the loss between the final output of the model *Ô*^(*t*)^ and actual target output *O*^(*t*)^ which is defined as

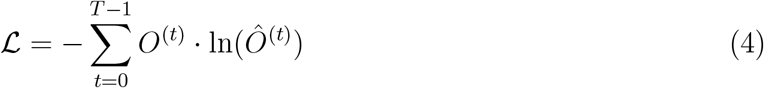

where *T* represents the total time of the trajectory, equivalent to the sequence length (*l*). The model has been trained over 10000 epochs, with all hyperparameters across all of the systems provided in the Table-S2. Figures S1(a-f) show the training and validation loss as a function of epochs for six distinct systems that would be elaborated in the upcoming sections of the present article. The plots indicate that the loss curves saturate or fluctuate slightly about the mean after a certain number of epochs, suggesting robust training of the transformer model without overfitting.

#### C. Generating future states from pre-trained transformer

After training the transformer model, it can generate any desired number of time series future states by inputting an initial sequence of tokens as a prompt. For a given sequence, the model will generate a probability distribution over the entire vocabulary/states. From this probability distribution, the next element of the sequence can be sampled using a multinomial distribution (see supplemental results SR1 for details). As we have generated the future states across all systems using our trained model, hereafter we will refer to it as the Generative Pre-Trained Transformer (GPT) model. The GPT model was built using PyTorch^18^ and our implementation is available in Github at the following URL: https://github.com/palash892/gpt_state_generation

### GPT precisely captures inter-state transition kinetics in model multi-state systems

To begin, we will delve into two hand-crafted model systems: 2D Brownian dynamics (BD) simulations of a single particle in 3-state and 4-state potentials. The mathematical representations of the potentials and simulation details are provided in the “Methods” section. We employ the BD trajectories to compute the 2D free energy of the two model systems within their *X* and *Y* coordinate space, defined as 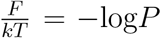, where *P* is the probability, calculated by using a 2D histogram of the coordinates. Figure 2(a) represents the free energy surface (FES) plot for the 3-state toy models. The plot exhibits three minima in the FES and the particle can hop from one minimum to another. The states are marked in the plots using magenta color, identified through K-means clustering in coordinate space (Figure 2(b)). After clustering the data, the entire trajectory is discretized into a specific number of states. Figure 2(c) shows the trajectory after spatial discretization, where each cluster index corresponds to a metastable state. The trajectory demonstrates that the particle can stay in a particular state for some time, and also exhibits transitions between various states. Now, from both the actual (i.e. BD-simulated) and GPT-generated time series data, we can compute the probability of each state by counting the occurrences of that particular state and dividing by the total count. For instance, if the count of state 0 is *C*_0_ and the total count of all states is *C*_*tot*_, then the probability of state 0 is 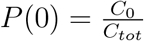. Figure 2(d) depicts a comparison between the actual and GPT-generated state probabilities for the 3-state model. The plot suggests that there is a close match between the actual and GPT-generated state probabilities.

**Figure 2.**
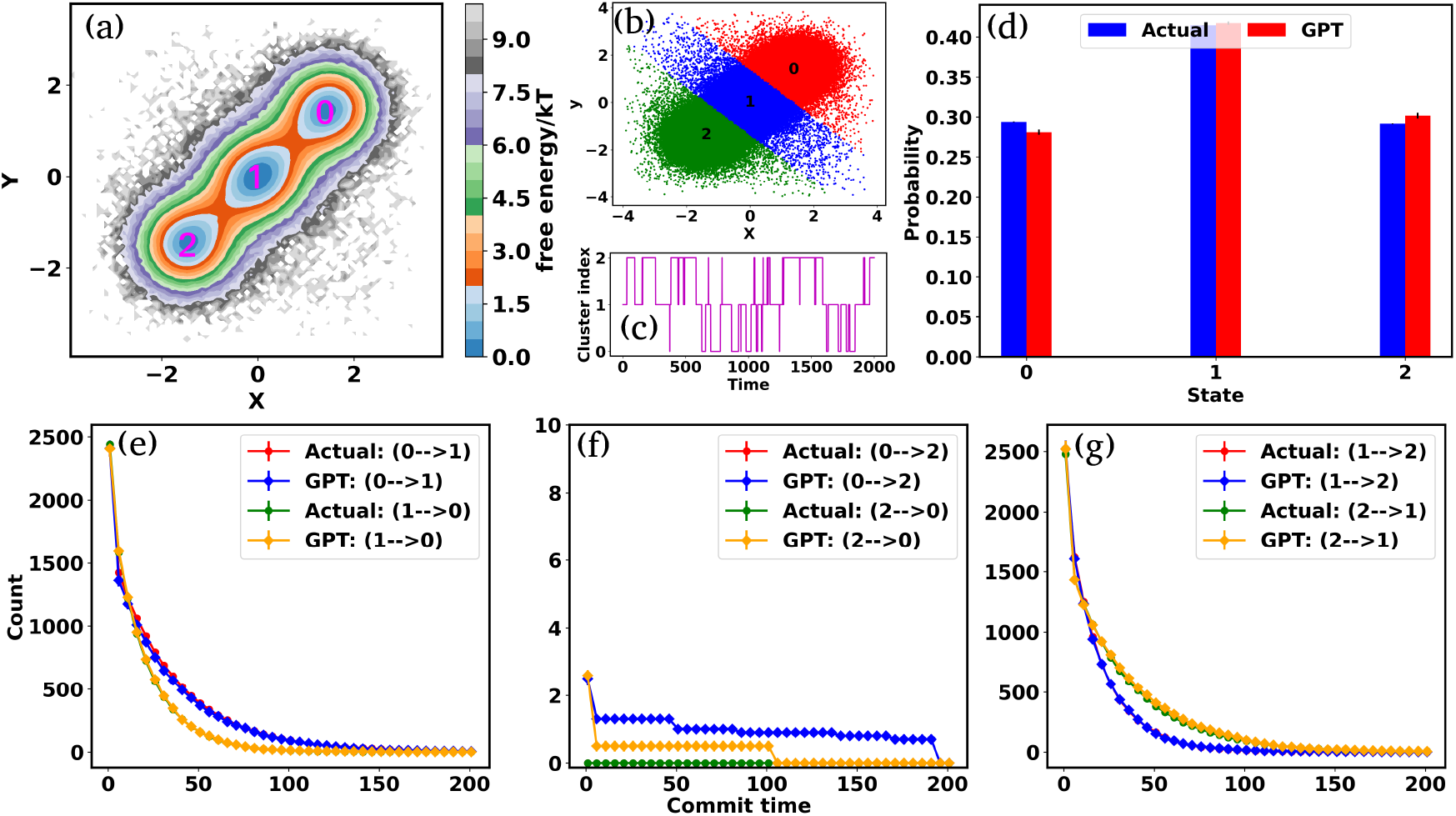
Kinetics and thermodynamics for toy model system. (a) Free energy surface (FES) plot for 3-state toy model in their *X* and *Y* coordinate space. The particle can transition from one minimum to the other. (b) Scatter plots of the *X* and *Y* coordinate, with distinct clusters representing metastable states identified through K-means clustering. (c) The trajectory of the particle in 3-state potential after state decomposition. (d) The comparison of state probabilities between the actual and GPT-generated time series data for the 3-state toy model. The plot highlights the accuracy of the GPT model in predicting the state probabilities. (e-g) Transition counts as a function of commit time for a 3-state toy model. These plots indicate the ability of the GPT model to learn contextual relationships among the states and generate future states that are kinetically and thermodynamically significant. Here the error bar represents the standard error and the commit time is in the unit of *τ*_*BD*_ (see “Methods”).

To effectively compare the kinetics between actual and GPT-generated time series data, one must analyze the transitions between different states over time. To facilitate this analysis, we utilized the concept of “commitment time or commit time”. This metric represents the duration a particle remains in a given state before transitioning to another. ^16,17^ We calculated the total transition count for different commit times, considering all possible state pairs (^*n*^*C*_2_,*n* is the total number of states) and both forward and backward directions. Figures 2(e-g) represent the transition count as a function of commit time for a 3-state toy model. These plots reveal that the decay of transition counts as a function of commit time is very similar between actual and GPT-generated data in both directions. Furthermore, the 2D FES plot (Figure 2(a)) indicates that states 0 and 2 are spatially distant. The actual data corroborates this by showing no direct transitions between these states. Quite interestingly, the GPT model captures a maximum of three transitions, showcasing its capability to capture the underlying dynamics.

We assessed the accuracy of this approach for a 4-state toy model system, as depicted in Figures S2(a-j)., Barring slight deviations in probabilities and a few transitions (Figure S2(f) and Figure S2(i)), the results are very similar for both actual and GPT-generated states. The free energy surface (FES) plot in Figure S2(a) indicates that states 1 and 0, as well as states 2 and 3, are spatially distant, with no direct transitions between them in the actual data. Remarkably, the GPT-generated states capture these trends very nicely. However, upon closer examination, the FES (Figure S2(a)) suggests that although states 0 and 2, and states 1 and 3, are spatially proximate, the trajectory is not sampled properly. This discrepancy may contribute to the slight deviations in transition counting between actual and GPT-generated data for these pair of states (0-2, 2-0, 3-1, and 1-3). Nevertheless, the results collectively indicate that the GPT model effectively learns the context and relationships between states, enabling it to generate future states that are both kinetically and thermodynamically significant.

### Predicting the ensemble probabilities and state-to-state transition kinetics in Trp-cage mini protein

Encouraged by the promising results in model potential, we extended our approach on a globular 20-residue mini-protein Trp-cage. This biomolecule is known for complex and multi-state conformational ensemble, despite its small size and remains an ideal candidate for experimental and computational studies of protein folding. Towards this end, we utilized a long (100*µs*) unbiased simulation trajectory of Trp-cage, provided by D. E. Shaw Research.^19,20^ For this system, due to higher degrees of freedom, it is necessary to define suitable order parameters or collective variables (CVs) to discretize the time series data into a certain number of states. Nonetheless, the identification of precise CVs is a challenging problem. To address these challenges, we employ an encoder-decoder-based unsupervised deep neural network called Autoencoder.^21,22^ An Autoencoder is a powerful non-linear dimension reduction technique that is used to transform the high-dimensional input data in a lower-dimensional space known as the latent space. The encoder component of the Autoencoder maps the input data to the latent space, while the decoder reverses this process, reconstructing the original input from the latent space. During this process, the model optimizes its weights and biases to preserve the most important information from the input data in the lower-dimensional representation. Figure 3(a) represents a schematic of autoencoder architecture where the distances between *C*_*α*_ atoms serve as input features. During the training of the Autoen-coder, we monitor the fraction of variation explained (FVE) score to determine the optimal dimension of the latent space. A detailed description of the Autoencoder architecture and the various hyperparameters are provided in the (“Methods” section and Table-S3).

**Figure 3.**
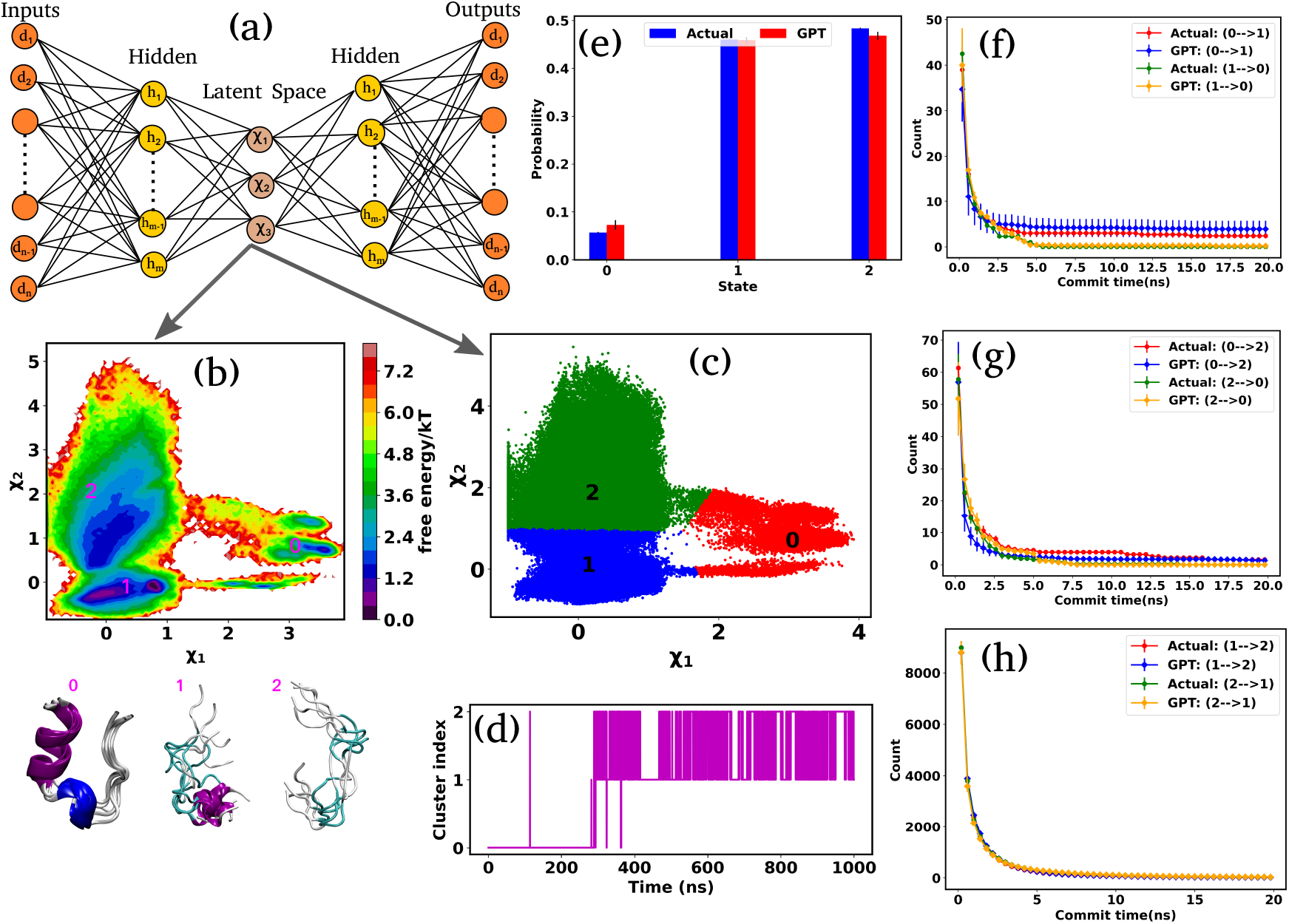
Kinetics and thermodynamics for Trp-cage mini protein. (a) A schematic representation of the Autoencoder. In this setup, *d*_1_, *d*_2_, …, *d*_*n*_ denote the input and output nodes, delineating the dimensions of the input and output data, while *h*_1_, *h*_2_, …, *h*_*n*_ represent the hidden nodes. Furthermore, *χ*_1_, …, *χ*_3_ represent the latent nodes. (b) 2D FES plot along latent space *χ*_1_ and *χ*_2_ for Trp-cage with three distinct minima and extracted conformations. (c) The state decomposition of the MD trajectory is achieved through k-means clustering on the latent space, which divides the entire trajectory into distinct sequences of states. (d) The trajectory of the Trp-cage after state decomposition. (e) The comparison of state probabilities for between actual and GPT-generated time series data. These plots suggest that the GPT model effectively captures the probabilities with minor deviations. (f-h) Comparison of transition counts between actual and GPT-generated states, showcasing the GPT model’s ability to accurately capture state transitions. Here error bar represents the standard error.

Figure 3(b) represents the 2D FES plot along the latent space *χ*_1_ and *χ*_2_, obtained from Autoencoder. The figure clearly shows three distinct minima in the FES plot, indicating distinct conformations. To visualize different conformations, we extracted a few conformations near each minimum in the FES and overlaid them. The superimposed conformations reveal mainly three metastable states: folded *α*-helix (state-0), partially folded *α*-helix (state-1), and unfolded random coils (state-2). After clustering in the latent space, the entire trajectory comprises metastable states and their transitions, as depicted in Figure 3(c). Figure 3(d) illustrates the discretized trajectory, with the majority of transitions occurring between states 1 and 2. Figure 3(e) represents a comparison of the state probabilities between the actual and GPT-generated time series data for the Trp-cage protein. These figures demonstrate that the GPT model has effectively captured the probabilities, with minor deviations observed for a few states. Importantly, these deviations are within the error bars.

Next, to probe the kinetics between various states of the Trp-cage protein, we calculated the transition counts as a function of commit time, akin to methodologies employed for 3-state and 4-state toy models. Figures 3(f-h) compare the transition counts between actual and GPT-generated states for the Trp-cage protein. These figures encompass all possible pairs and transitions in both forward and reverse directions. The results indicate that the GPT model accurately captures the transitions between states. Quite interestingly, the GPT model can also predict the highest number of transitions between states 1 and 2 accurately which aligns with our observations from the trajectory itself (Figure 3(e)). In summary, together these results suggest that the GPT model effectively captures the probability and the transition counts between various states of the Trp-cage protein, providing valuable insights into the kinetics and thermodynamics of these systems.

### On intrinsically disordered protein *α*-Synuclein

Next, we turn our attention to an intricately complex system: *α*-Synuclein. It is a prototypical intrinsically disordered protein (IDP) that predominantly resides in the human brain, especially nerve cells.^23^ It plays a crucial role in the communication between these cells by releasing neurotransmitters which are messengers chemicals. ^24^ However, the excessive accumulation of these proteins can lead to several neurodegenerative disorders such as Parkinson’s disease and other synucleinopathies.^25–27^ For *α*-Synuclein, we utilize the 73*µs* trajectory provided by D. E. Shaw Research (DESRES).^19,20^

We used the radius of gyration (*R*_*g*_) as CVs to decompose the entire trajectory into a specific number of states. Figure 4(a) represents the (*R*_*g*_) (red color) of the *α*-Synuclein as a function of time, showcasing the protein’s diverse conformational possibilities. Through K-means clustering in the *R*_*g*_ space,^28^ we discretize the total trajectories into distinct states (Figure 4(a) magenta color plot). Specifically, three states have been identified: intermediated compact (state-0), collapsed (state-1), and extended (state-2), with a superimposition of snapshots revealing significant conformational heterogeneity within each state (Figures 4(b)).

**Figure 4.**
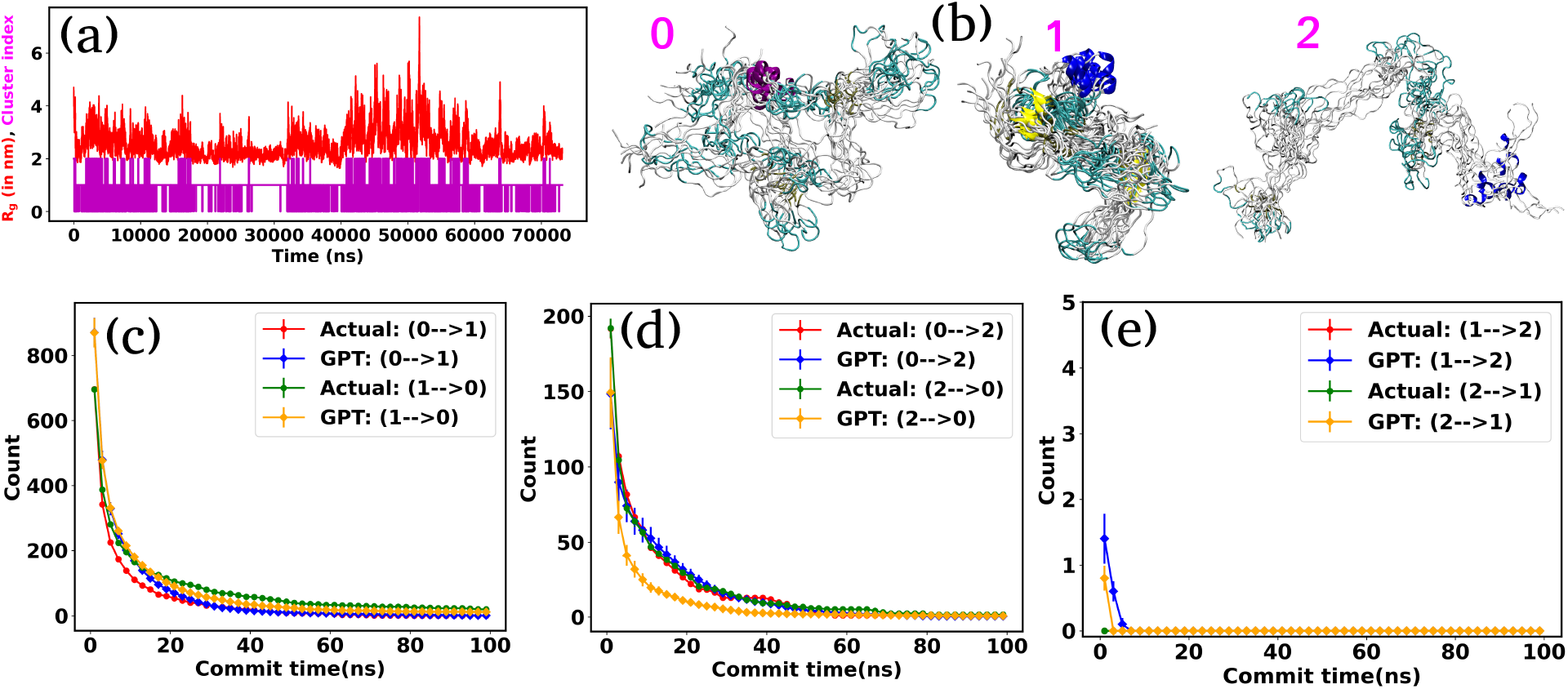
The radius of gyration (*R*_*g*_) and transition count comparison for *α*-Synuclein. (a) The (*R*_*g*_) as a function of time for *α*-Synuclein (red plot). The magenta color plot represents the trajectory after state decomposition via K-means clustering. (b) The diverse conformational states are identified as intermediated compact (state-0), collapsed (state-1), and extended (state-2). (c-e) Transition count comparison between the actual and GPT-generated time series data as a function of commit time for *α*-Synuclein. The transition dynamics fairly match with actual and GPT-generated data except at a smaller commit time. Here the error bar represents the standard error.

Figure S3 compares the state probabilities between actual and GPT-generated time series data for *α*-Synuclein. The figure clearly shows the deviation in the state probability values. Next, to analyze the transition dynamics, we have calculated the transition count of each state. Figures 4(d-f) depict the comparison of transition count as a function of commit time between actual and GPT-generated time series data for *α*-Synuclein. While there are some deviations in probability values, the transition dynamics align fairly well with actual and GPT-generated data, except for a smaller commit time. Specifically, the GPT model generates a lower count for transition (2 → 0) compared to actual data. The deviations in state probability and transition dynamics indicate the intricacies involved in accurately predicting the dynamical behavior of such a complex system.

### Assessing transformer’s predictive ability in far-from-equilibrium system: active worm-like polymer chain

In the preceding sections, we have mainly focused on capturing kinetics and thermodynamics of various models and real systems that are in thermodynamic equilibrium. Finally, in this section, we shift our attention to an active system. Most living organisms are active and their movement is powered by energy consumption from internal or external sources. The continuous consumption and dissipation of energy drive these systems far from equilibrium. Notably, the activity or self-propulsion force plays a crucial role in the formation of many self-organized collective phenomena such as pattern formation,^29–31^ motility-induced phase separation,^32–34^ swarming motion,^35–38^ etc. In this study, we employ a model system, an active worm-like polymer chain,^39^ where the activity or self-propulsion force acts tangentially along all bonds. We have utilized our in-house BD simulation trajectory to study the active polymer chain (see Methods for details).

After training the Autoencoder by using inter-bead distances as features, we have chosen two-dimensional latent space as CVs. Figure 5(a) shows the FES plot across the latent space *χ*_1_ and *χ*_2_ for the active polymer chain and the corresponding metastable states are highlighted in magenta color. The overlaid plots suggest that there are mainly two metastable states: a bending state (state-1) and a spiral state (state-0). Notably, despite the apparent simplicity of the two states, the visualization of the trajectory (Figure S4(a)) suggests that the system spends a very long time within each state along with spontaneous spiral formation and breakup occurrences. Subsequently, we employ K-means clustering on the latent space derived from Autoencoder to discretize the trajectory (Figure 5(b)).

**Figure 5.**
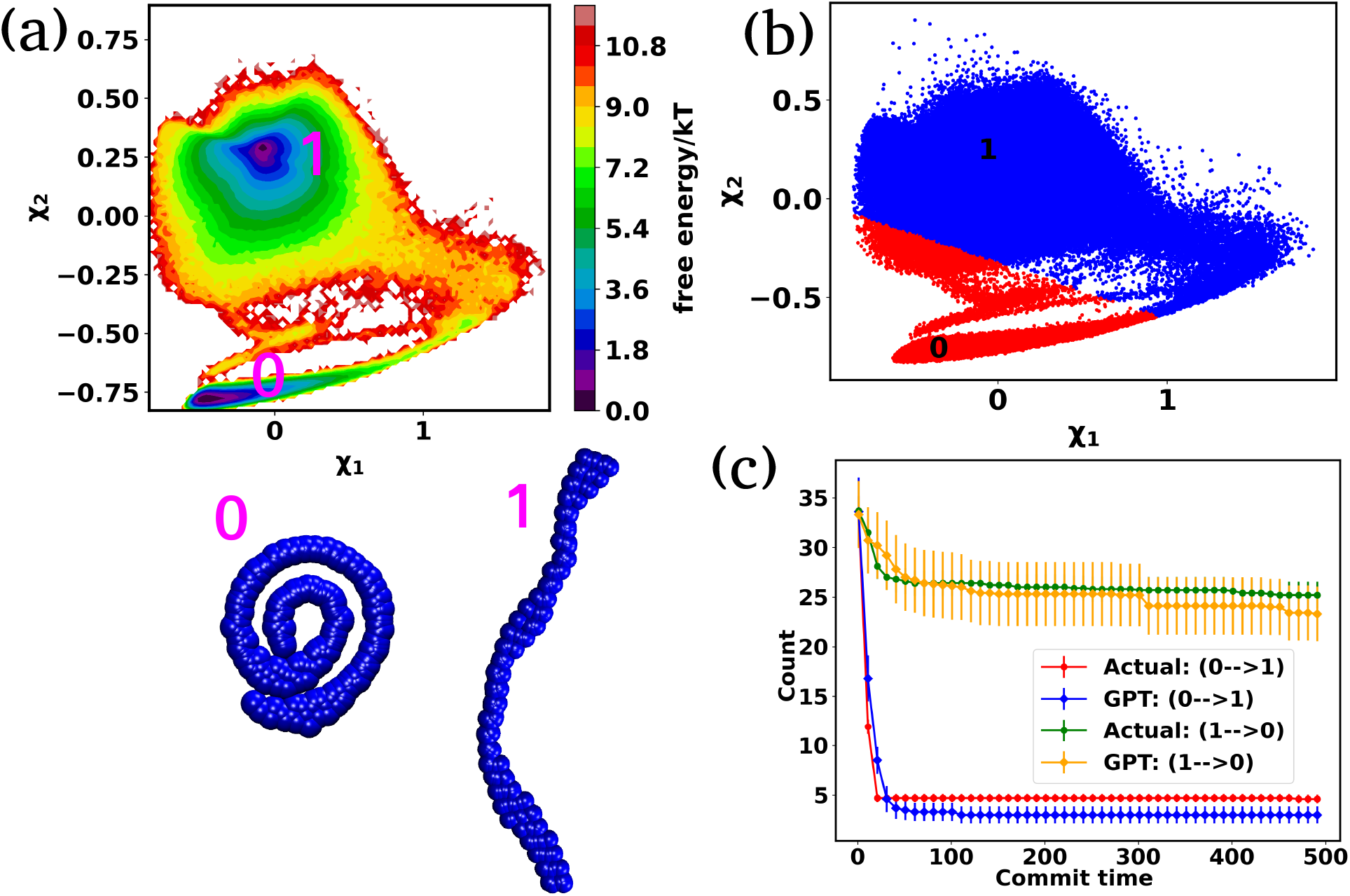
The free energy surface and transition count comparison for the active worm-like polymer chain. (a) The FES plot of the active polymer chain across the latent space *χ*_1_ and *χ*_2_. This plot highlights two metastable states: a bending state (state-1) and a spiral state (state-0). (b) The latent space has been clustered via K-means clustering to discretize the trajectory. (c) The comparison of transition count between actual and GPT-generated data for the active polymer chain, reflecting the system’s long stay at particular states before transition and the violation of detailed balance. The GPT model accurately generates future states, maintaining saturation and detailed balance violation, albeit with some small deviation from actual data. Here the error bar represents the standard error and the commit time is in the unit of *τ*_*BD*_ (see “Methods”).

Figure S4(b) compares the state probabilities between actual and GPT-generated time series data for active polymer chain. The comparison reveals the deviations in the state probability. Figure 5(c) depicts the comparison of transition count as a function of commit time between actual and GPT-generated time series data. However, interestingly, the GPT model accurately generates the transition for the active system. As mentioned earlier, for the active polymer chain, the system stays at a particular state for a long time before transit to the other state. This behaviour is reflected in the plots, where both curves saturate after a certain commit time. Moreover, the forward and backward transition curves suggest a violation of detailed balance, which is an inherent feature of active systems that are far from thermodynamic equilibrium. Remarkably, the GPT model successfully generates future states that maintain the saturation nature as well as the violation of detailed balance, albeit with some deviation from actual data within the error bar. These findings indicate that the GPT model is very powerful for future state predictions, even in complex active systems. In a similar spirit, for comparison purpose, we have also computed similar metrics for a ‘passive’ polymer chain for an enhanced clarity (Figures S5). Here too, we observed a strong alignment between the kinetics and thermodynamics captured by the GPT model and the actual BD-generated data.

### Analysing the impact of the attention layer

In the previous sections, our focus has been on exploring the thermodynamics and kinetics of diverse model and real systems, whether in thermodynamic equilibrium or out of equilibrium. We have observed that despite variations in state probabilities, the GPT model generates future states that accurately maintain transition dynamics in a statistical sense. Now, in this section, we delve into identifying the pivotal factors that maintain these precise transition dynamics.

In the field of natural language processing (NLP), the seminal work by Vaswani et al. titled “Attention Is All You Need”^7^ introduces the paradigm-shifting concept of self-attention. The state-of-the-art transformer model, based on attention mechanisms, has demonstrated superior performance compared to traditional recurrent neural networks (RNNs). To understand the role of attention, we computed the attention score from the multi-head attention layer which is defined as 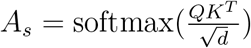 (see equation 3 for details), where *Q, K*, and *d* are the query, key, and the dimension of the embedding layer, respectively. In our study, we randomly selected 20 chunks of sequence length 128 from GPT-generated time series data and fed them as inputs for the trained GPT model. Subsequently, we computed the attention scores from each head, averaging them across all heads and the 20 random chunks. Physically, these attention scores unveil correlations among different tokens within a sequence of time series data. Figures 6(a-f) show the heat map of the masked attention score for all the systems. These plots highlight the presence of significant finite, non-zero attention among various tokens within sequences. Notably, some tokens exhibit clear evidence of long-range attention, underscoring the model’s ability to understand the relationships between events that are far apart.

**Figure 6.**
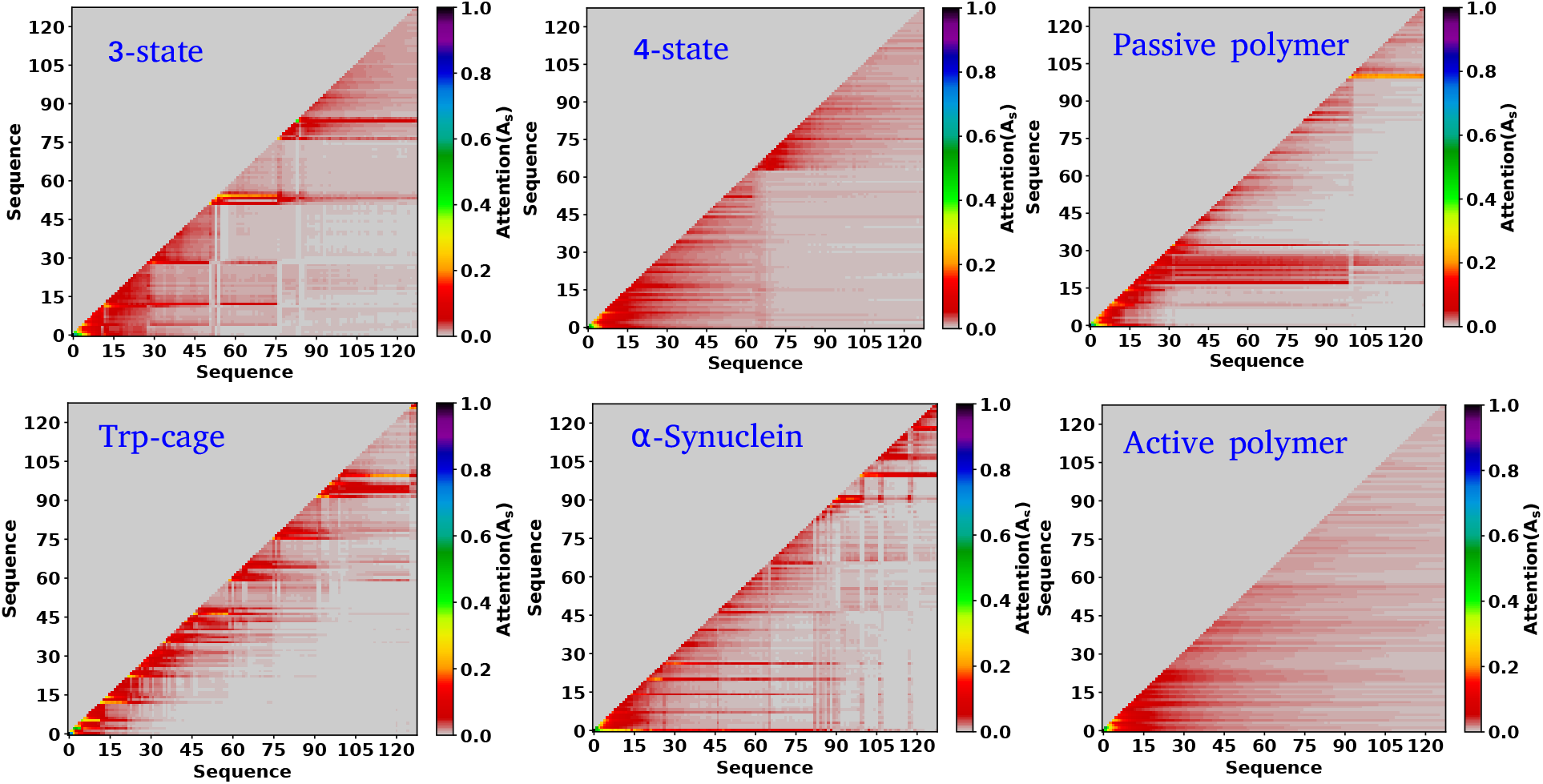
Attention score for all the systems analyzed. (a-f) The heat map of the attention score computed from the multi-head attention layer of a trained GPT model, for six distinct systems under consideration. These scores reveal that there is finite attention among various tokens and highlight the evidence of long-range attention.

Based on our examination of attention scores, we now pose the following question: How does attention impact the transition dynamics between different states? To address this, we trained the GPT model on all six systems by removing the multi-head attention layer, while maintaining other hyperparameters consistent with our previous model architecture. This approach ensures that the model can no longer capture any short or long-range correlations between tokens. Our findings indicate that for simple systems, such as a 3-state toy model, the transition dynamics captured by the GPT model remain similar regardless of the presence or absence of the attention layer (see Figure S6). This suggests that attention may not play a significant role in these simple systems. However, for other systems, there are substantial deviations in GPT-generated transition dynamics compared to actual data. Figure 7(a-j) represents the comparison of transition count as a function of commit time between the actual and GPT-generated time series data for all of the systems under consideration as highlighted in the text of each plot. These plots suggest that while in many cases the GPT model can recover the transition counts in one direction, it completely predicts the wrong transitions in other directions. Here, we have only shown the plots where the deviations are prominent. The comprehensive plots for all the systems with all possible transitions are given in SI (Figures S7-S10). Together these analyses identify the power of the attention layer. Even in physicochemical systems, attention is crucial for learning the context and relationships or correlations among various tokens of time series data, enabling the model to generate a physically meaningful future state.

**Figure 7.**
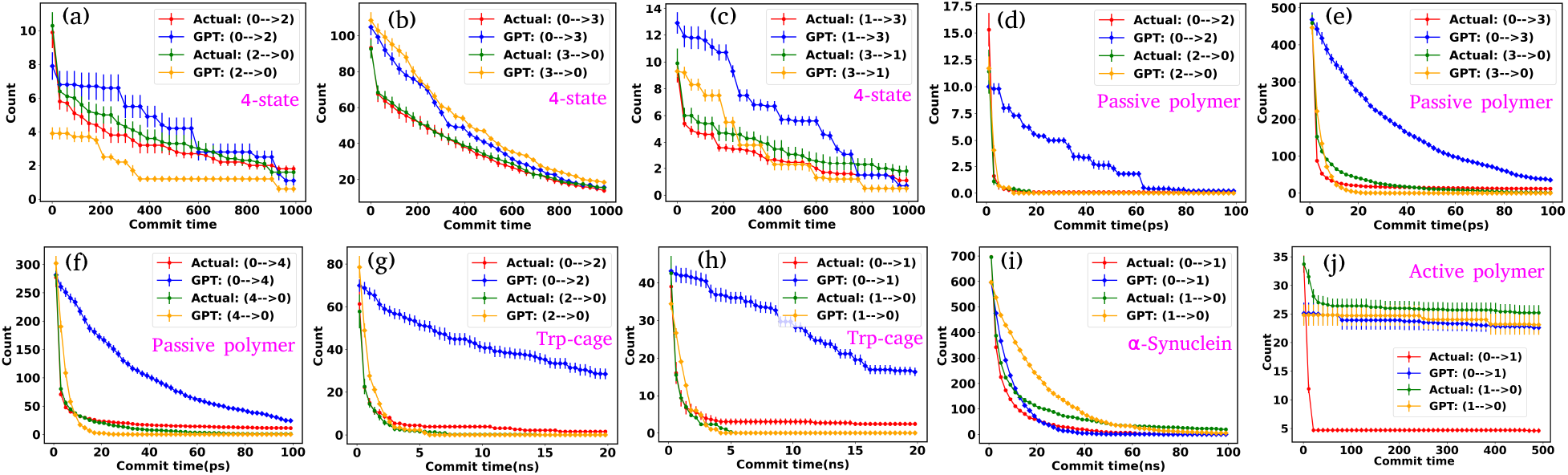
Impact of Attention on Transition Dynamics in GPT Model. (a-j) The comparison of transition counts over commit time between the actual and GPT-generated time series data for all of the systems in absence of attention mechanisms. There are significant deviations in GPT-generated transition dynamics compared to the actual data. While the GPT model can sometimes accurately predict transitions in one direction, it frequently mispredicts transitions in another direction.

### Performance Comparison: MD Simulation vs. GPT State Generation

After training the GPT model, the generation of the future state is extremely fast compared to conventional MD simulations of equivalent systems. For example, the model can generate 20000 next sequence for *α*-Synuclein within ∼ 6.0 minutes, corresponding to a 20 *µs* simulation of this system with a data saving frequency of 1 ns. Similarly, the model can generate 100000 sequences for the Trp-cage mini protein within ∼ 32 minutes, equivalent to 20 *µs* simulation with a data dumping frequency of 200 ps. As mentioned earlier, we have analyzed six systems among them the Trp-cage mini protein and *α*-Synuclein are particularly relevant in biophysical contexts, and their simulations were conducted in real-time units. Thus, our primary focus here is to compare the performance of these two systems. To compare this generation efficiency against actual MD simulation times, we conducted 10 ns simulations for these systems, maintaining all parameters such as box size, salt concentration, temperature, and time step as per the study by Robustelli et al.^19^ As the data saving frequency for these two systems is different, we define a 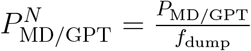 where *P*_MD/GPT_ is the performance of MD or GPT model and *f*_dump_ is the saving frequency of the data. This metric can normalize the performance of each system by its respective data saving frequency. All the MD simulations and training of the GPT models were performed on an Intel(R) Xeon(R) Platinum 8352Y CPU at 2.20GHz, along with an NVIDIA RTX A6000 GPU. Table 1 represents all of the details of the performance of the MD simulations as well as GPT state generation. The table 1 suggests that the performance and normalized performance of the GPT model outstrip those of traditional MD simulations, demonstrating its efficiency in generating future states of the systems.

**Table 1:** The comparison of the performance between MD simulation and GPT state generation.

| System | $f_{\text{dump}}$ | $P_{\text{MD}}$<br>(ns/day) | $P_{\text{MD}}^N$<br>(/day) | Training<br>time<br>(minutes) | Gen.<br>sequence | Gen.<br>time<br>(minutes) | $P_{\text{GPT}}$<br>($\mu\text{s}/\text{day}$) | $P_{\text{GPT}}^N$<br>($10^3/\text{day}$) |
| --- | --- | --- | --- | --- | --- | --- | --- | --- |
| α-Synuclein | 1.0 ns | $\sim 35.0$ | $\sim 35.0$ | $\sim 127.0$ | 20,000 | $\sim 6.0$ | $\sim 4800.0$ | $\sim 4800.0$ |
| Trp-cage | 200 ps | $\sim 832.0$ | $\sim 4160.0$ | $\sim 35.0$ | 100,000 | $\sim 32.0$ | $\sim 900.0$ | $\sim 4500.0$ |

What should be the typical size of training data? To evaluate the impact of training data size on our model’s ability to predict future states in a biophysical system, we trained our GPT model using varying amounts of data. We selected the Trp-cage mini protein, which provides 500, 000 frames, in contrast to the 73, 124 frames available for *α*-Synuclein (see Table S1). We generated the same number of states as depicted in Figure 3 but varied the training data size. Initially, the model was trained with 60% of the total data. We have now conducted additional training with 10%, 40%, and 50% of the data. Figures S11(a-i) and S11(j-l) show the transition counts over time and state probabilities for these different training data percentages. These figures indicate a significant alignment between the Actual and GPT-generated data after utilizing 40% of the training data. This suggests that a sufficient amount of training data is crucial for the model to predict future states accurately. However, the transition counts plots demonstrate that even with just 10% of the training data, the model can still capture complex relationships between various states, generating future states that are not entirely erroneous.

## Discussions

The time evolution of biophysical systems undergoes various conformation changes, depending on environmental conditions. Understanding these dynamics typically requires experiments or Molecular Dynamics (MD) simulations. However, comprehending the long-term behaviour of these systems requires running computationally expensive simulations. In this study, we present a comprehensive approach, employing state-of-the-art machine learning models, particularly decoder-only transformers to predict the future states of physicochemical systems. Through an extensive analysis of the MD trajectory of various models and real systems, we have demonstrated the efficacy of the GPT model in capturing both the kinetics and thermodynamics of these systems.

Our study began with simplified model systems, namely 3-state and 4-state toy models. We employed K-means clustering on coordinate space for this simple system to discretize the trajectory. Subsequently, we delved into more complex systems such as the Trp-cage mini protein, 32-bead passive polymer chain, and intrinsically disordered protein *α*-Synuclein. However, for these systems, we utilized another ML-based technique Autoencoder, to identify the relevant collective variables for discretization. Our results highlight the ability of the GPT model to accurately predict the probabilities of different states and capture the transition dynamics between them. Furthermore, we extended our analysis to include an active system, a 32-bead active worm-like polymer chain, where the system is far from thermodynamic equilibrium. Remarkably, the GPT model successfully predicted the kinetics and thermodynamics of the active system.

An interesting aspect of our study is to dissect the role of the attention mechanism within the transformer architecture. Through attention score analysis, we noticed that there are significant correlations among different tokens within sequences, indicating the model’s capacity to understand long-range dependencies. Furthermore, by removing the attention layer from the model, we observed substantial deviations in transition dynamics for complex systems. Therefore, these findings strongly suggest that the attention mechanism plays a pivotal role in maintaining accurate predictions.

Although the transformer-based large language models (LLM) were specially developed for tasks like machine translation and natural language processing, our study demonstrates their effectiveness in predicting the kinetics and thermodynamics of a diverse array of biophysical systems. One notable limitation of our GPT model is that it never generates new states beyond its training data. In terms of language, the transformer-based model is always unable to generate new vocabulary. Nonetheless, the model can learn the complex syntactic and semantic relationships present in the sequence of tokens which help them to generate future states of the system very correctly. If some system shows a completely new state at a very long time that is not present in the training sequence, the model can not generate that particular state. This suggests that one needs very good MD sampling data for a physical system to predict the future state of the system. However, recent advances in ML have introduced innovative techniques such as variational autoencoders (VAE), ^40–42^ generative adversarial networks (GANs),^43,44^ diffusion models,^45,46^ etc., which can be utilized for generating new conformations and improving sampling in biophysical systems. ^47–49^ While these techniques enhance sampling quality, they may lack the temporal information crucial for understanding the dynamics of the system. In conclusion, our findings highlight the potential of LLMs as powerful tools for advancing our understanding of biophysical systems, offering new avenues for further exploration and improvement in this field.

## Methods

### 3-state and 4-state toy model

To simulate a particle in a 2D 3-state and 4-state potential well, we adopted the same functional form for the potential as Tsai et al.^16^ The potential for the 3-state model is given by:

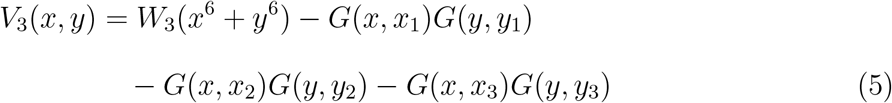

Similarly, the potential for the 4-state model is given by:

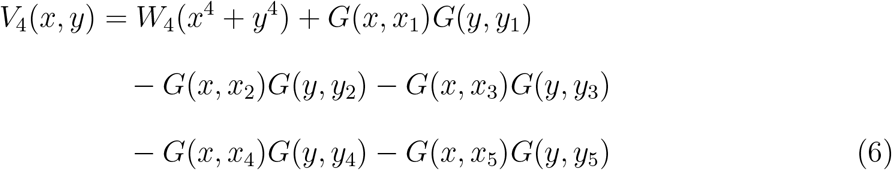

Where *G* is a Gaussian function as 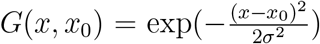. Here *x*_0_ and *σ* are the mean and standard deviation of the distribution. In our simulation we kept *W*_3_ = *W*_4_ = 0.0001 and *σ* = 0.8. The mean values of Gaussian distribution for the 3-state model are given by *x*_1_ = 0.0, *y*_1_ = 0.0, *x*_2_ = *−*1.5, *y*_2_ = *−*1.5, and *x*_3_ = 1.5, *y*_3_ = 1.5. Similarly, for the 4-state model, these values are *x*_1_ = 0.0, *y*_1_ = 0.0, *x*_2_ = 2.0, *y*_2_ = *−*1.0, *x*_3_ = 0.5, *y*_3_ = 2.0, *x*_4_ = *−*0.5, *y*_4_ = *−*2.0, and *x*_5_ = *−*2.0, *y*_5_ = 1.0. We performed Brownian dynamics simulations for these two systems by integrating the equation of motion:

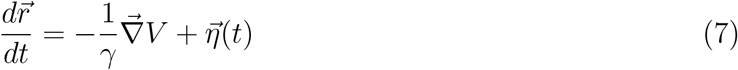

where *γ* is the friction coefficient, *V* is the potential energy, and 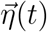 is a random noise, satisfying fluctuation dissipation theorem i.e 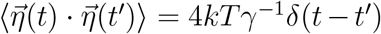. Here all simulations time is in the unit of 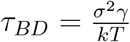, where *σ* = 1 is the diameter of the particle. Integration of the equation 7 was performed using Euler method with a time step *δt* = 0.01*τ*_*BD*_ by setting 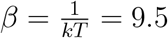.

### Passive polymer chain

We performed a very long molecular dynamics simulation (5.74*µs*) for a passive polymer chain, with the model and simulation parameters of the polymer system detailed in a previous study by our group.^50^

### Trp-cage mini protein and *α*-Synuclein

For Trp-cage and *α*-Synuclein, we utilized very long molecular dynamics simulation trajectories from D. E. Shaw Research. ^19^ The Trp-cage trajectory spanned 100*µs*, while the *α*-Synuclein trajectory was 73*µs* long. These simulations were performed using the a99SB-disp force field on Anton specialized hardware.^20^ The detailed simulation protocols can be found in the original paper by Robustelli et al.^19^ However, in the original paper, the trajectory of *α*-Synuclein was 30*µs* long. Due to some periodic image issues in the original trajectory, the authors provided the extended 73*µs* trajectory for *α*-Synuclein, maintaining the same simulation setup as before but increasing the box size.

### Active worm-like polymer chain

The two-dimensional worm-like polymer chain consists of *N* = 32 beads connected by a stiff spring with spring constant *k*_0_. Each bead has a diameter of *σ* and an equilibrium bond length of *d*_0_. The dynamics of the polymer chain are governed by overdamped motion, described by the equation:

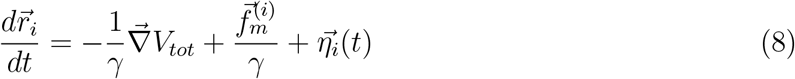

where *γ* is the friction coefficient, and *V*_*tot*_ is the total potential energy of the system, which includes bonding potential *V*_*bond*_, bending potential *V*_*bend*_, and non-bonded potential *V*_*nb*_. The bonding potential *V*_*bond*_ is given by:

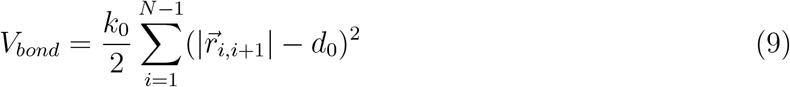

The bending potential *V*_*bend*_ is given by:

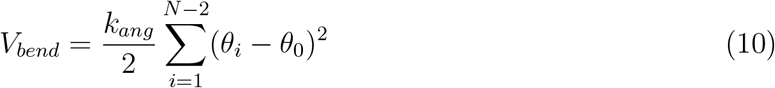

where *k*_*ang*_ is the bending rigidity and *θ*_0_ is the equilibrium bond angle. The non-bonded potential *V*_*nb*_ is taken as the Hertzian potential:

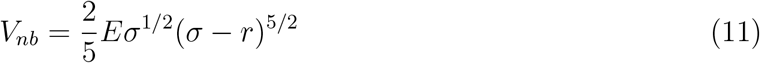

These potentials are very generic for the simulation of any passive polymer chain. However, the most important force for an active polymer chain is the motility or self-propulsion force. Here, 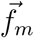 represents the self-propulsion force which acts tangentially along all bonds. The random noise 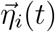 maintains the fluctuation-dissipation theorem i.e 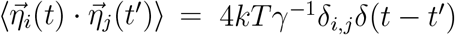. All lengths and energies are in the units of *σ* and *kT*, respectively, and the simulation time is in the units of 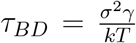. The Brownian dynamics simulation for an active polymer chain was performed using a time step of *dt* = 0.001*τ*_*BD*_. The other simulation parameters are *kT* = 1.0, *σ* = 1.0, *d*_0_ = 0.5*σ*, 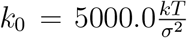, *k*_*ang*_ = 45.0*kT*, *γ* = 200*kTτ*_*BD*_*/σ*^2^, *f*_*m*_ = 5.0*kT/σ, θ*_0_ = *π*, and *E* = 10000.0*kT/σ*^3^

### Training details of Autoencoder

The Autoencoder architecture and training hyperparameters are presented in Table-S3. For the Trp-cage mini protein, we utilized the distance between all *C*_*α*_ atoms as input features, resulting in 190 (^20^*C*_2_ = 190) features for its 20 residues. Conversely, for active and passive polymer chains, we employed inter-beads distances, selecting 8 effective beads in an arithmetic progression with a step of 4, providing 28 (^8^*C*_2_ = 28) input features.^51^ Throughout the Autoencoder training process, we monitored two metrics: training loss and the fraction of variation explained (FVE) score. The FVE is defined by the equation:

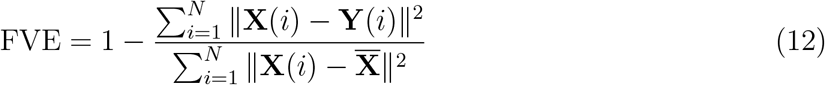

where, **X**(*i*), **Y**(*i*), and 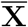 represent the input, output, and mean input, respectively, and *N* corresponds to the total number of features. The FVE score indicates the proportion of input data variance explained by the Autoencoder’s reconstruction. Figures S12(a-f) depict the plots of these two metrics for all systems as described within the figure text. For active and passive polymer chains, a 2D latent dimension was chosen, whereas for the Trp-cage mini protein, a 4D latent dimension was utilized. Across all these latent dimensions, the FVE scores exceed 0.80, indicating that the Autoencoder’s reconstruction explains at least 80% of the original data variance. Furthermore, the gradual decrease followed by saturation of the training loss curve suggests no overfitting during the Autoencoder’s training process. However, in the case of the IDP *α*-Synuclein, even with a high latent dimension of *L*_*d*_ = 7, we have observed a relatively low FVE score (*∼* 0.60) (Figure S13(a)). Additionally, we have plotted the FES using the first two components of the latent space (Figure S13(b)). This plot indicates a lack of distinct minima in the free energy surface. Consequently, clustering the data for a proper state decomposition within this latent space is not feasible. Therefore, for *α*-Synuclein, we have opted to use the radius of gyration (*R*_*g*_) as the reaction coordinate to discretize the trajectory into a specific number of states.

We conducted simulations for 3-state, 4-state, and active worm-like polymer chains using our in-house scripts written in C++. To simulate passive polymer chains, we utilized the open-source Software GROMACS-2022.^52,53^ Our Autoencoder model was trained using Python implementation of Tensorflow^54^ and Keras^55^ and the GPT model was built using PyTorch.^18^

## Data and code availability

The manuscript contains all the data. The code and detailed documentation for training the Autoencoder and GPT model are available on GitHub at the following URL: https://github.com/palash892/gpt_state_generation

## Acknowledgments

All the authors acknowledge Tata Institute of Fundamental Research Hyderabad, India for providing the support of computing resources. We acknowledge support of the Department of Atomic Energy, Government of India, under Project Identification No. RTI 4007. JM acknowledges Core Research grants provided by the Department of Science and Technology (DST) of India (CRG/2023/001426). We would like to thank D. E. Shaw Research for providing us with access to a long simulation trajectory of Trp-cage and monomeric *α*-Synuclein.

## Notes

### Competing Interest Statement

The authors have declared no competing interest.

### Summary of Updates

A new analysis and supplemental figures were introduced.

