## supplemental methods, tables and figures for "Predicting Future Kinetic States of Physicochemical Systems Using Generative Pre-trained Transformer"

**SR1:**Generation of the next state of a sequence using multinomial distribution:
Let's assume for an event, we have $k$ classes ($k$ possible outcomes) and $n$ number of trials. For example, tossing a coin has two possible outcomes, and throwing a die has six possible outcomes. Now $x_1, x_2, ...x_k$ are the frequencies of each outcome, such that $\sum_{i=1}^{k} x_i = n$ and $p_1, p_2, ...p_k$ are the probabilities of each outcome occurring in a single trial, such that $\sum_{i=1}^{k} p_i = 1$. Then the probability distribution function is given by:

$$P(x_1, x_2, ...x_k) = \frac{n!}{x_1!x_2! \cdots x_k!} \cdot p_1^{x_1} p_2^{x_2} ... p_k^{x_k} \tag{1}$$

Binomial distribution is the special case of multinomial distribution where $x_1 + x_2 = n$, and $p_1 + p_2 = 1$, then the distribution will be

$$P(x_1) = \frac{n!}{x_1!(n - x_1)!} \cdot p_1^{x_1} p_2^{n-x_1} \tag{2}$$

As discussed in the main text for a given sequence, the GPT model will generate a probability distribution over the entire vocabulary/states. From this probability distribution, the next element of the sequence can be sampled using a multinomial distribution. Let's consider a four-state model, where the probability distribution of the next sequence is gen-prob = $[0.3286, 0.2406, 0.1770, 0.2539]$. These four values represent the probabilities of the four states, which are 0, 1, 2, and 3, respectively. The generation of the next state in the sequence is achieved by passing these probability values through a multinomial distribution. The multinomial distribution maintains these probability values and uses them to determine the next state in the sequence.

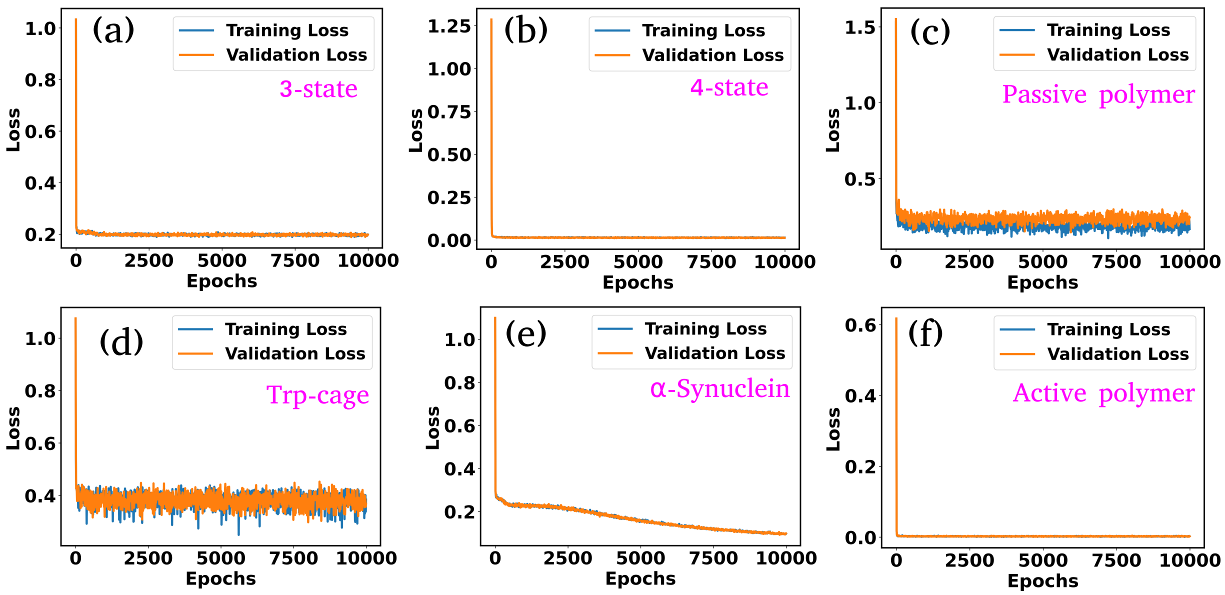


FIG. S1. Training and validation loss as a function of epochs for all of the system under investigation

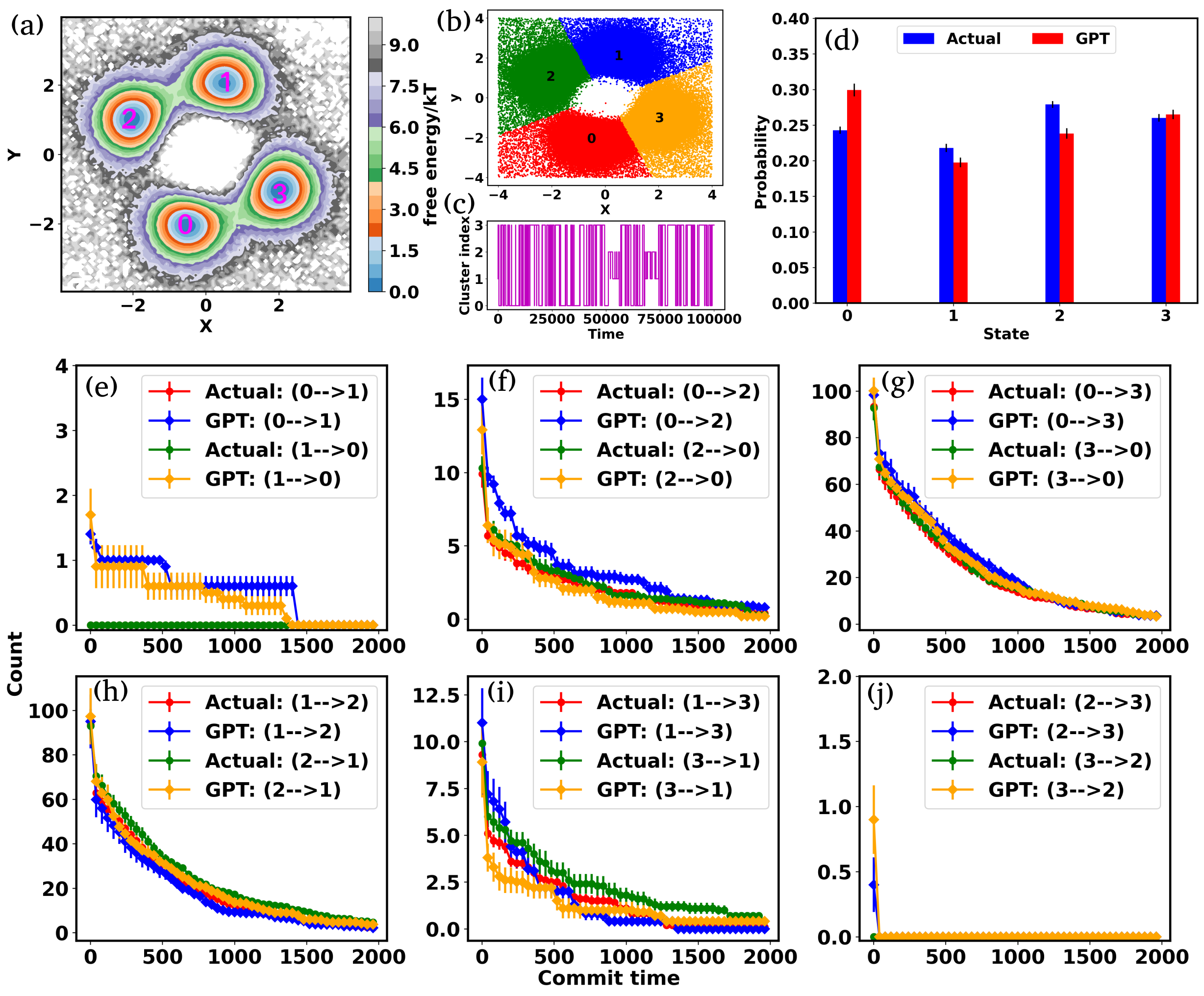


FIG. S2. (a) Free energy surface (FES) plot for 4-state toy model in their $X$ and $Y$ coordinate space. The particle can transition from one minimum to the other. (b) Scatter plots of the $X$ and $Y$ coordinate, with distinct clusters representing metastable states identified through K-means clustering. (c) The trajectory of the particle in 4-state potential after state decomposition. (d) The comparison of state probabilities between the actual and GPT-generated time series data for the 4-state toy model. (e-j) Transition counts as a function of commit time for a 4-state toy model. Here the error bar represents the standard error

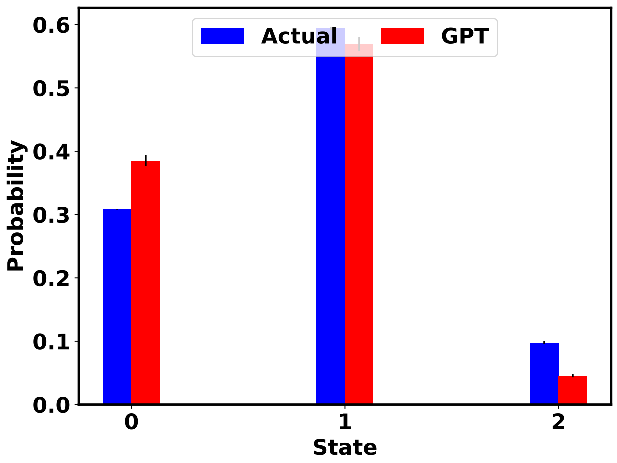


FIG. S3. The comparison of state probabilities between the actual and GPT-generated time series data for IDP $\alpha$-Synuclein.

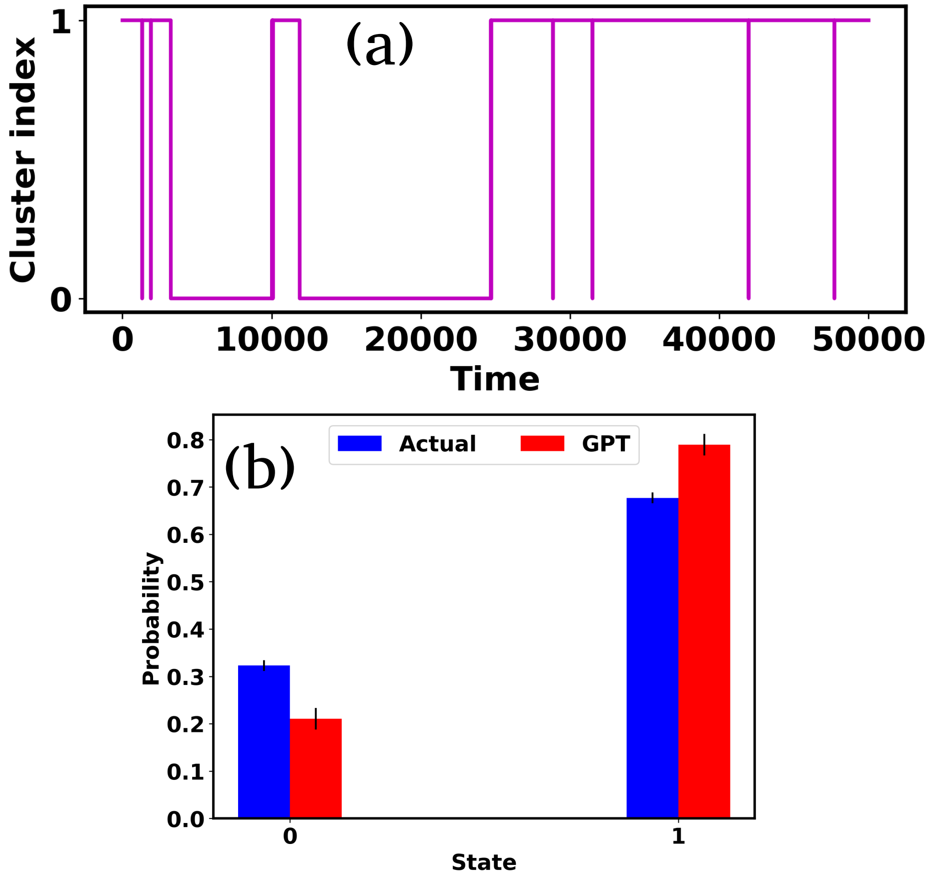


FIG. S4. (a) The trajectory of active worm like polymer chain after state decomposition. (b) The comparison of state probabilities between the actual and GPT-generated time series data for active worm like polymer chain.

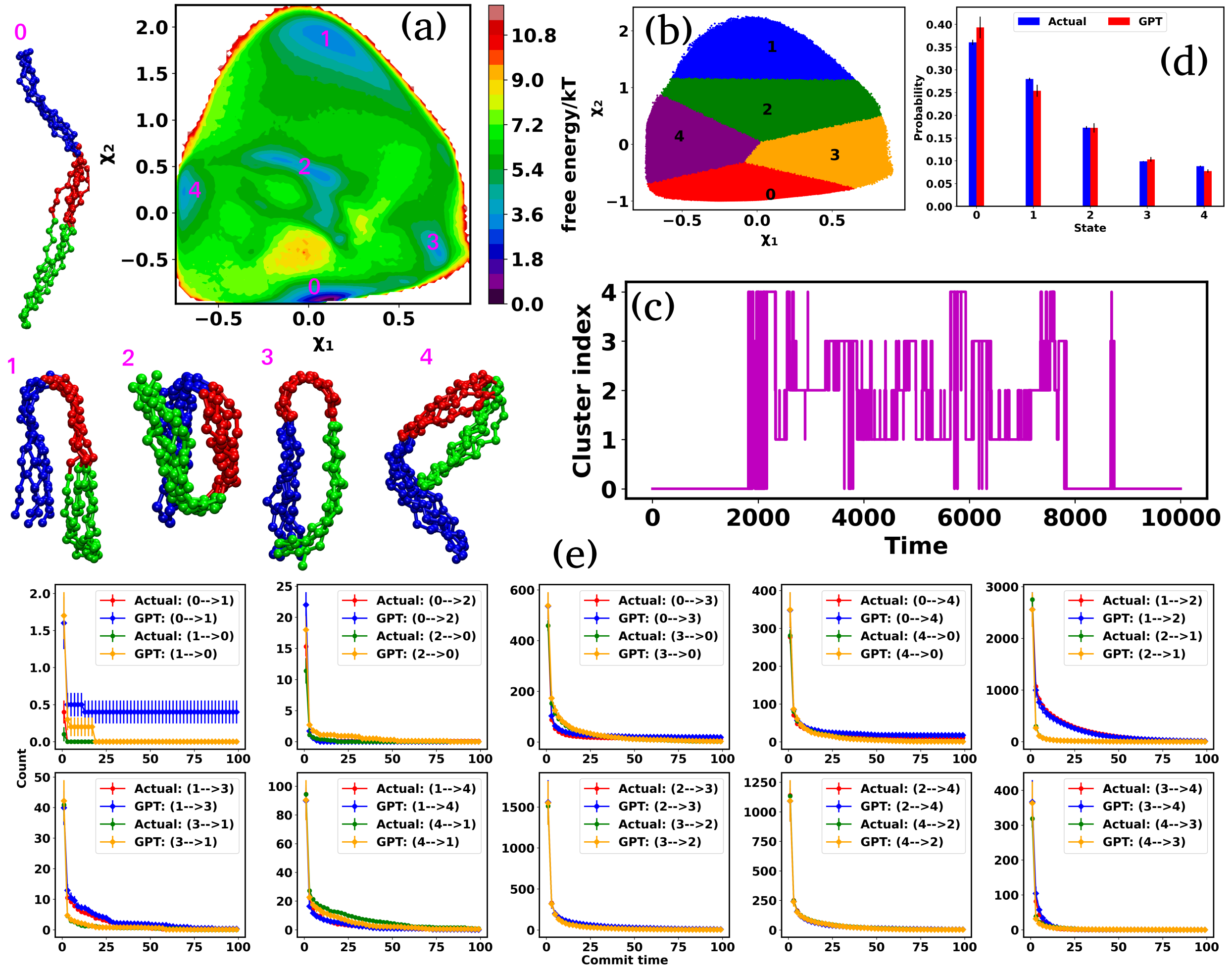


FIG. S5. (a) (a) 2D FES plot along latent space $\chi_1$ and $\chi_2$ for passive polymer chain, showing five distinct minima and extracted conformations. (b) The state decomposition of the MD trajectory is achieved through k-means clustering on the latent space. (c) The trajectory of the passive polymer chain after state decomposition. (d) The comparison of state probabilities between the actual and GPT-generated time series data for passive polymer chain. (e) Comparison of transition counts between actual and GPT-generated states for the passive polymer chain, showcasing the GPT model's ability to accurately capture state transitions. Here error bar represents the standard error.

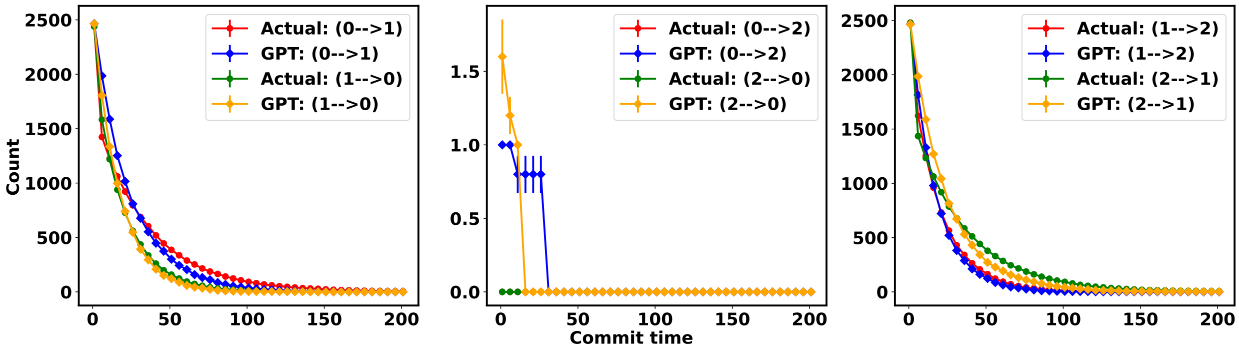


FIG. S6. The comparison of transition counts over commit time between the actual and GPT-generated time series data for 3-state toy model potential in absence of attention mechanism.

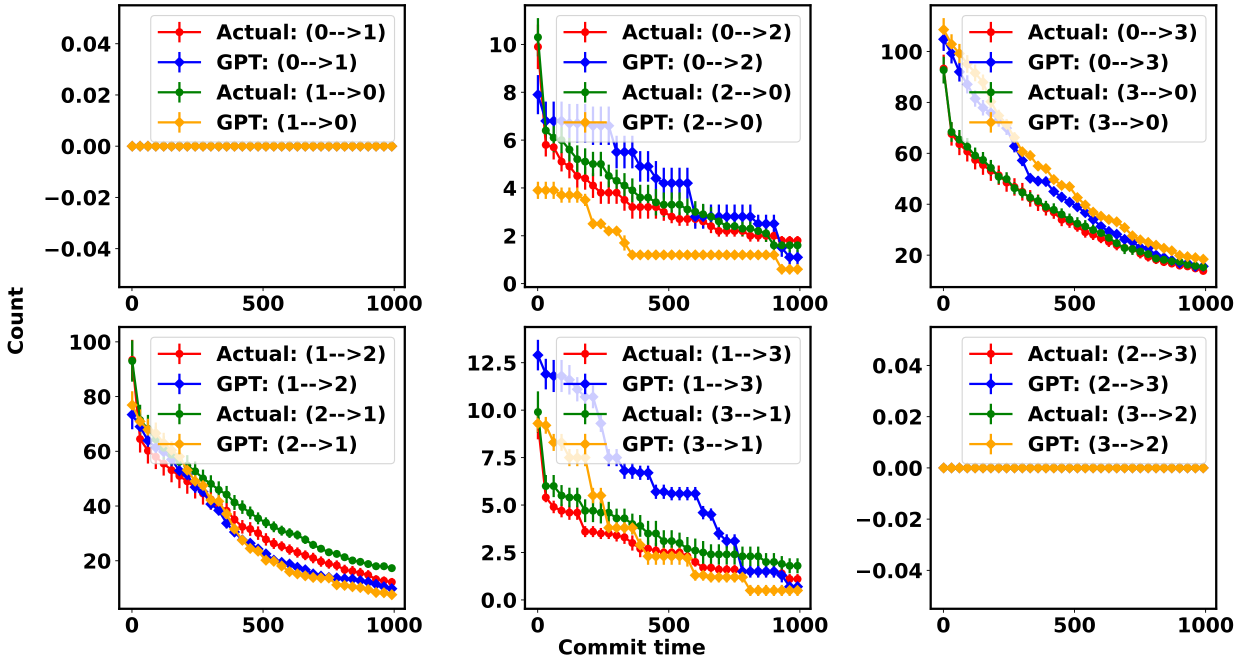


FIG. S7. The comparison of transition counts over commit time between the actual and GPT-generated time series data for 4-state toy model potential in absence of attention mechanism.

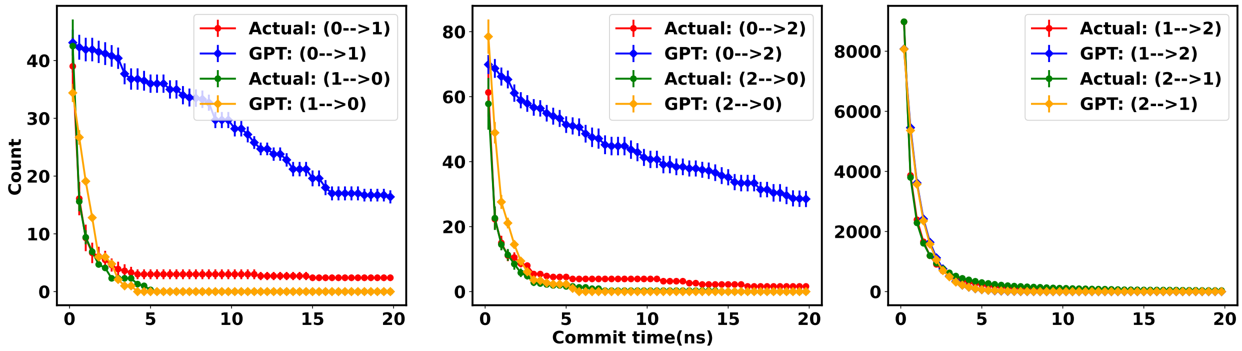


FIG. S8. The comparison of transition counts over commit time between the actual and GPT-generated time series data for Trp-cage mini protein in absence of attention mechanism.

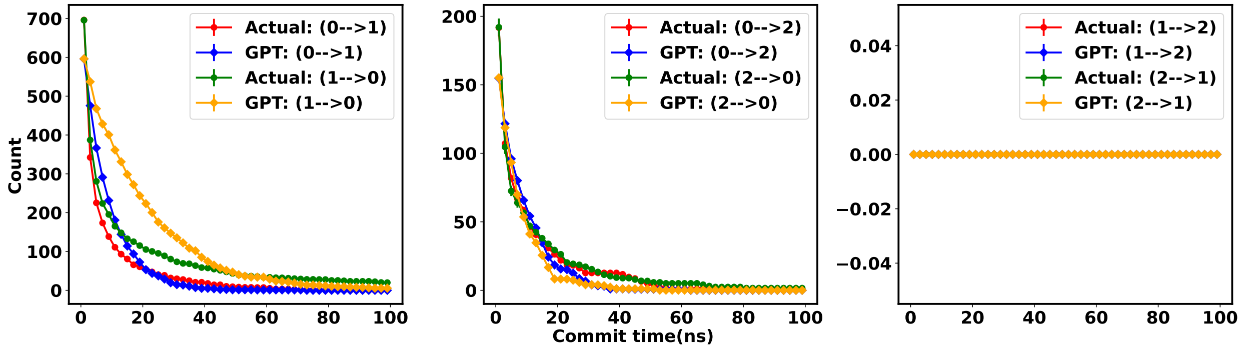


FIG. S9. The comparison of transition counts over commit time between the actual and GPT-generated time series data for $\alpha$-Synuclein in absence of attention mechanism.

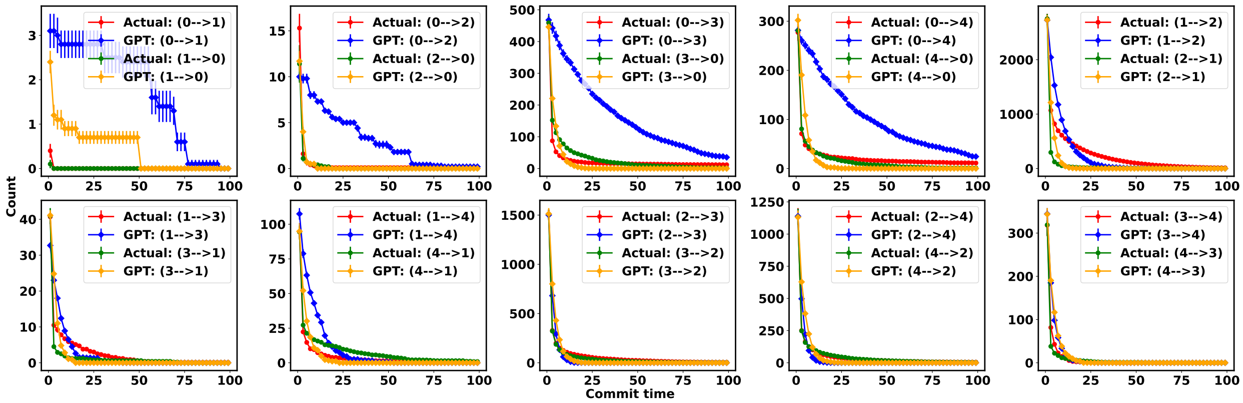


FIG. S10. The comparison of transition counts over commit time between the actual and GPT-generated time series data for passive polymer chain in absence of attention mechanism.

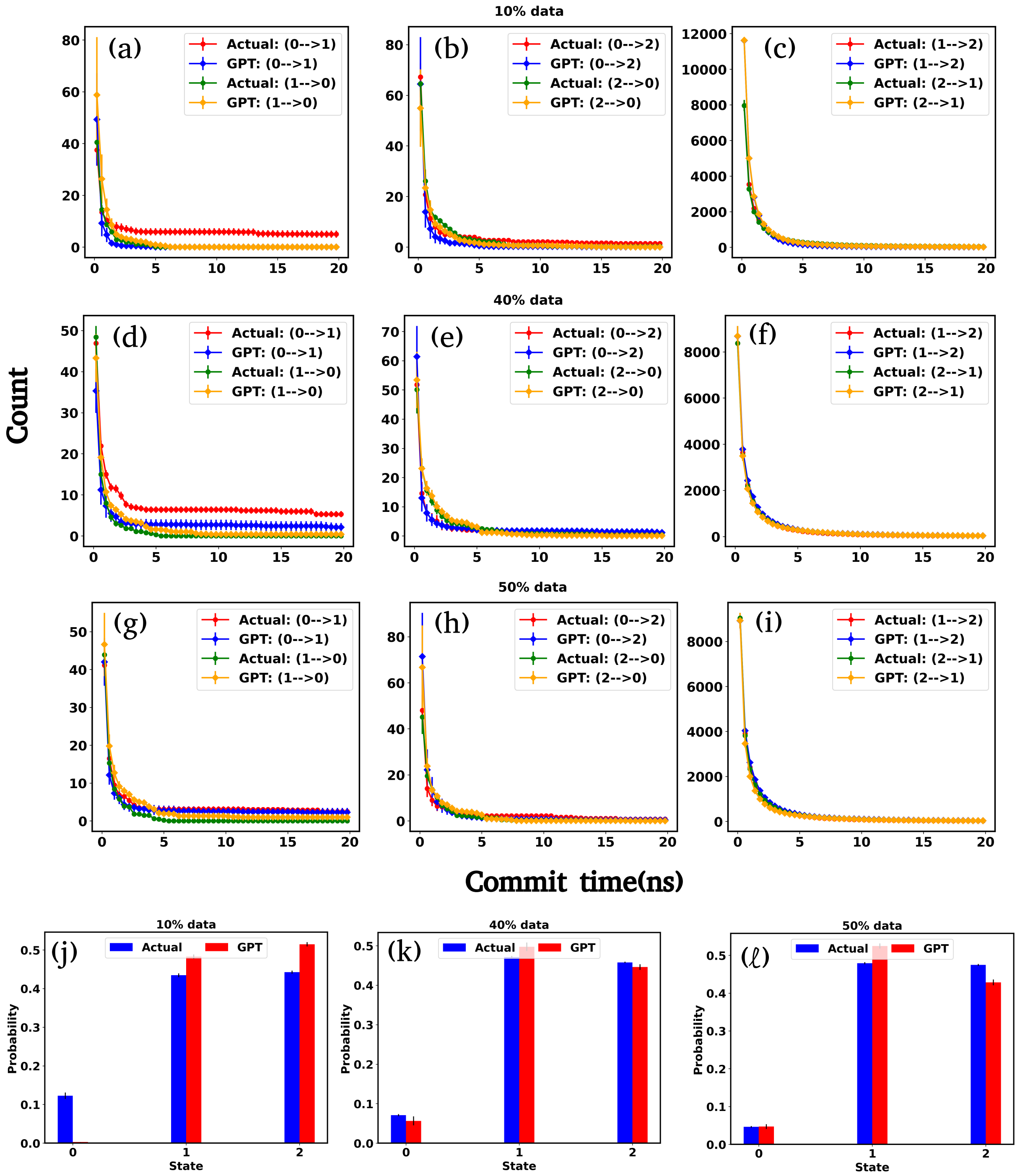


FIG. S11. (a-i) The comparison of transition counts between actual and GPT-generated states for various percentage of training data of Trp-cage. (j-l) The comparison of state probabilities between the actual and GPT-generated time series data for various percentage of training data of Trp-cage.

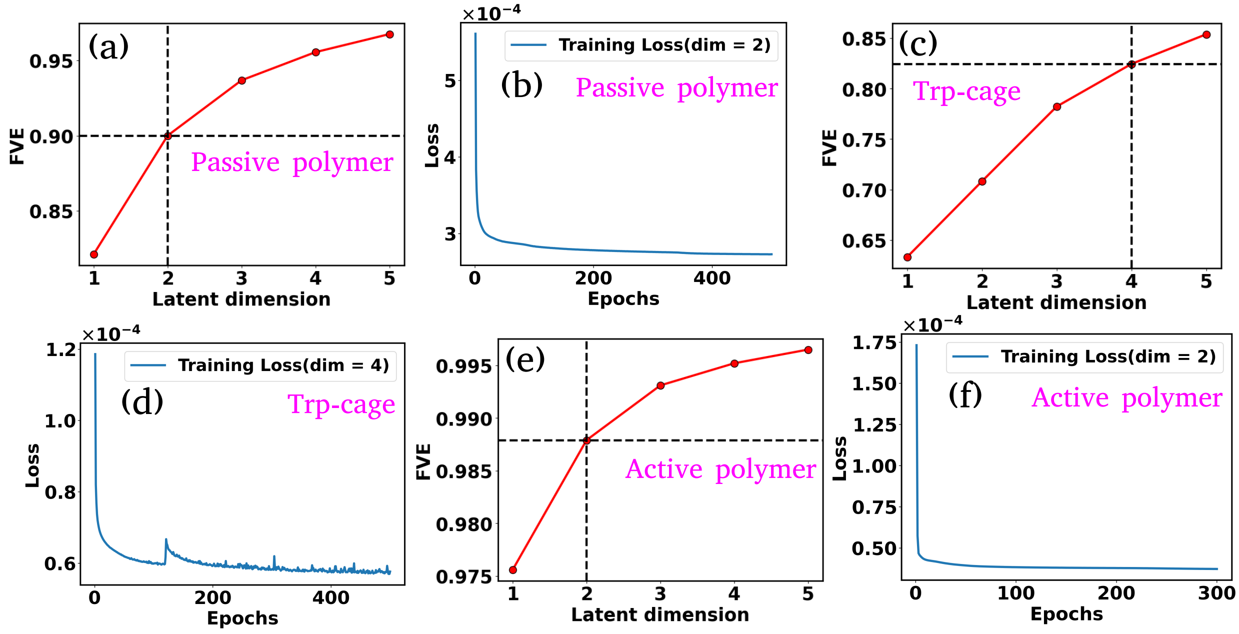


FIG. S12. (a-f) FVE and training loss plots for all systems. We have chosen 2D latent dimensions for active and passive polymer chains and 4D latent dimensions for Trp-cage mini protein.

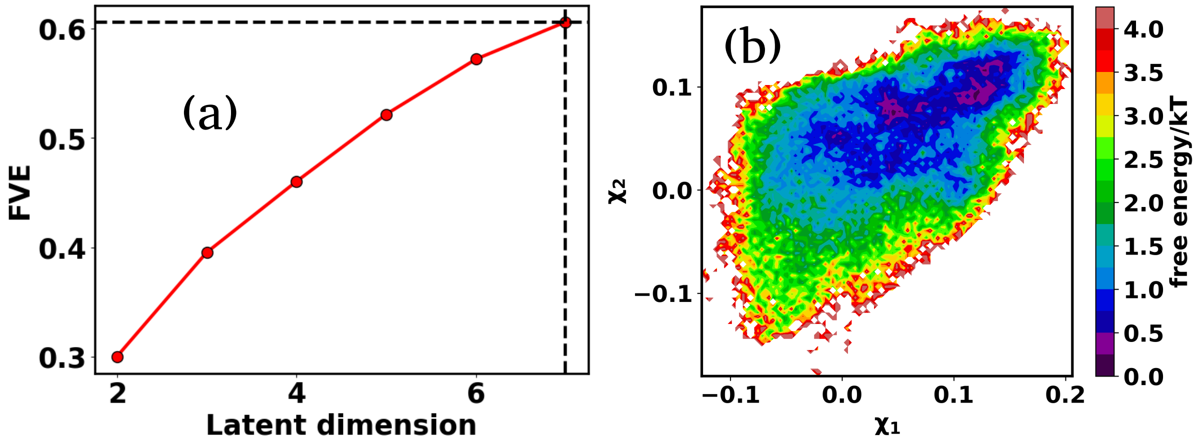


FIG. S13. (a) The FVE score as a function of latent dimension for $\alpha$-Synuclein. The value of the FVE score is approximately 0.60, even with a high latent dimension of $L_d = 7$. (b) The FES plot along the first two latent dimensions for $\alpha$-Synuclein reveals no distinct minima. As a result, state decomposition through clustering within this latent space is unfeasible.

TABLE S1. The amount of data used for training, validation and testing

| System | Total data | Training | Validation | Prompt | Testing or generation |
|---|---|---|---|---|---|
| 3-state | 50,00000 | 18,00000 | 2,00000 | 128 | 2,00000 |
| 4-state | 50,00000 | 18,00000 | 2,00000 | 128 | 2,00000 |
| Passive polymer | 57,47973 | 18,00000 | 2,00000 | 128 | 2,00000 |
| Active polymer | 50,00000 | 18,00000 | 2,00000 | 128 | 2,00000 |
| Trp-cage | 5,00000 | 2,70000 | 30,000 | 128 | 1,00000 |
| $\alpha$-Synuclein | 73,124 | 50,000 | 5,000 | 128 | 20,000 |

TABLE S2. The hyperparameters used for training the GPT model.

| System | batch size | block size | epochs | embeddings($d$) | head($N_h$) | blocks($N_b$) | learning rate | optimizer |
|---|---|---|---|---|---|---|---|---|
| 3-state | 128 | 128 | 10000 | 256 | 8 | 8 | $10^{-4}$ | AdamW [1] |
| 4-state | 128 | 128 | 10000 | 256 | 8 | 8 | $10^{-4}$ | AdamW |
| Passive polymer | 128 | 128 | 10000 | 256 | 8 | 8 | $10^{-4}$ | AdamW |
| Active polymer | 128 | 128 | 10000 | 256 | 8 | 8 | $10^{-4}$ | AdamW |
| Trp-cage | 128 | 128 | 10000 | 256 | 8 | 8 | $10^{-4}$ | AdamW |
| $\alpha$-Synuclein | 128 | 384 | 10000 | 256 | 8 | 8 | $10^{-4}$ | AdamW |

TABLE S3. The hyperparameters used for training the Autoencoder. Here $L_d$ represents the latent dimensions

| System | batch size | epochs | Activation | learning rate | optimizer | loss function | architecture |
|---|---|---|---|---|---|---|---|
| Passive polymer | 100 | 500 | Elu | $5 \times 10^{-4}$ | Adam [2] | MSE | 28-12-$L_d$-12-28 |
| Active polymer | 100 | 300 | Elu | $5 \times 10^{-4}$ | Adam | MSE | 28-12-$L_d$-12-28 |
| Trp-cage | 64 | 500 | Elu | $5 \times 10^{-4}$ | Adam | MSE | 190-72-36-12-$L_d$ -12-36-72-190 |
| $\alpha$-Synuclein | 64 | 100 | Tanh | $10^{-4}$ | Adam | MSE | 9730-4096-1024-512-128-16-$L_d$ -16-128-512-1024-4096-9730 |

[1] Ilya Loshchilov and Frank Hutter. Decoupled weight decay regularization. *arXiv preprint arXiv:1711.05101*, 2017.

[2] Diederik P Kingma and Jimmy Ba. Adam: A method for stochastic optimization. *arXiv preprint arXiv:1412.6980*, 2014.
